# Real-time tracking of complex ubiquitination cascades using a fluorescent confocal on-bead assay

**DOI:** 10.1101/268706

**Authors:** Joanna Koszela, Nhan T Pham, David Evans, Stefan Mann, Irene Perez-Pi, Steven Shave, Derek F J Ceccarelli, Frank Sicheri, Mike Tyers, Manfred Auer

## Abstract

The ubiquitin-proteasome system (UPS) controls the stability, localization and/or activity of the proteome. However, the identification and characterization of complex individual ubiquitination cascades and their modulators remains a challenge. Here, we report a broadly applicable, multiplexed, miniaturized on-bead technique for real-time monitoring of various ubiquitination-related enzymatic activities. The assay, termed UPS-confocal fluorescence nanoscanning (UPS-CONA), employs a substrate of interest immobilized on a micro-bead and a fluorescently labelled ubiquitin which, upon enzymatic conjugation to the substrate, is quantitatively detected on the bead periphery by confocal imaging. UPS-CONA is suitable for studying individual enzymatic activities, including various E1, E2 and HECT-type E3 enzymes, and for monitoring multi-step reactions within ubiquitination cascades in a single experimental compartment. We demonstrate the power of the UPS-CONA technique by simultaneously following ubiquitin transfer from Ube1 through Ube2L3 to E6AP. We applied this multi-step setup to investigate the selectivity of five ubiquitination inhibitors reportedly targeting different classes of ubiquitination enzymes. Using UPS-CONA, we have identified a new activity of a small molecule E2 inhibitor, BAY 11-7082, and of a HECT E3 inhibitor, heclin, towards the Ube1 enzyme. As a sensitive, quantitative, flexible and reagent-efficient method with a straightforward protocol, UPS-CONA constitutes a powerful tool for interrogation of ubiquitination-related enzymatic pathways and their chemical modulators, and is readily scalable for large experiments.

## INTRODUCTION

Post-translational modification of protein substrates by the small modifier protein ubiquitin regulates the stability, localization, interactions and activity of a substantial fraction of the proteome. Disruption of the ubiquitin-proteasome system (UPS) is associated with many diseases, including cancer, neurological disorders, and immune system dysfunction ^1^, and as a consequence is a high priority area for drug discovery. In the canonical ubiquitination cascade, the C-terminal carboxylate of ubiquitin is first activated in an ATP-dependent fashion by an E1 ubiquitin-activating enzyme, followed by thioester transfer to an E2 ubiquitin-conjugating enzyme, before subsequent transfer via an E3 ubiquitin ligase to a substrate lysine residue and attachment through an isopeptide linkage. Repetition of this cascade enables a ubiquitin chain to be assembled on the substrate through conjugation of further ubiquitin moieties to one of the seven lysine residues on ubiquitin itself. Different ubiquitin chain linkages can specify different fates for the substrate. For example, K48-linked chains target substrates to the 26S proteasome for rapid degradation, while K63-linked chains can control the assembly of various protein complexes. Protein monoubiquitination also occurs frequently and can direct protein interactions, for example in intracellular vesicular trafficking. Various ubiquitin modifications can be read by dedicated reader domains and can be reversed through the action of specific deubiquitinating enzymes. The human genome encodes two E1 enzymes, 38 E2 enzymes, over 600 E3 enzymes, over 100 deubiquitinating enzymes, as well as various associated scaffolding subunits. The combinatorial promiscuity of ubiquitin system enzymes, including redundant substrate targeting by different E3s, has made the deconvolution of UPS-mediated control far more difficult than originally envisioned, with implications for UPS drug discovery. In addition, only a limited number of small molecule inhibitors of the UPS are available as mechanistic probes and most of these inhibitors are only poorly characterized for specificity ^2^. The dearth of reliable methods to assess molecular mode of action of chemical inhibitors with the necessary level of quantification, time resolution and specificity has hindered UPS drug discovery ^3–5^.

The confocal scanning technology (CONA) was originally developed to enable screening of combinatorial chemical libraries directly on the solid bead-based support, which in effect serves as a nanoscale assay compartment ^6–11^. The confocal sectioning of a monolayer of ~100 μm-sized beads imaged on a fluorescence microscope equipped with a scanning stage enables the high-throughput, highly sensitive detection of binding events between a bead-linked small molecule and a fluorescently labelled target protein in solution ^6,8^. Bead-based screening methods are miniaturized, versatile, highly sensitive, quantitative and fast, and have proven successful in identification of numerous small molecule and peptidomimetic binders and inhibitors against difficult protein targets, such as HuR, Importin beta and LFA-1 ^9–11^.

To address the need for a robust technique attuned to the multitude of complex UPS reactions, we developed a fluorescence-based on-bead confocal imaging method, termed UPS-confocal fluorescence nanoscanning (UPS-CONA), for quantitative characterization of dynamic ubiquitination reactions in real-time. We demonstrate suitability of UPS-CONA for studying E1, E2 and HECT E3 activities, both individually and in an integrated enzyme reaction cascade. The modularity and miniaturization of UPS-CONA, in combination with high sensitivity, provides a powerful tool for deciphering multi-step reactions by tracking the kinetics of ubiquitin transfer in real-time. In proof-of-concept experiments, we apply UPS-CONA to investigate the specific activity of selected ubiquitination inhibitors and uncover novel inhibitory activities.

## RESULTS AND DISCUSSION

### A confocal on-bead ubiquitination assay – the concept

To exploit the advantages of the CONA technique in the ubiquitination field, we developed UPS-CONA as a method for detection and characterization of complex ubiquitin enzyme-mediated reactions. In its new, redefined form, the assay employs a substrate or enzyme of interest, which is immobilized on a polymer micro-bead, and a fluorescently labelled ubiquitin or ubiquitin-like protein modifier (ULM) in solution (Fig. 1a). Upon ubiquitin conjugation to the on-bead substrate or enzyme, the fluorescence emission intensity of conjugates is detected by confocal imaging through the equatorial cross-sectional plane of the beads and observed as a fluorescent ring. The high sensitivity of on-bead assays without the need for cumbersome washing steps is achieved by the reduction of the background fluorescence intensity through removal of out-of-focus light as a result of confocal imaging. Consequently, the time-lapsed detection of fluorescent ubiquitin allows monitoring of the reaction in real time. Using low concentrations of reactants, both on-bead and in solution, allows the fluorescence emission intensity detected at the cross-section of the beads to be linearly proportional to the amount of enzymatic conjugate (Supporting Information Figure S1). After acquisition of confocal images of beads in a multi-well test plate, the ring intensity of hundreds to thousands of micro-beads forming a monolayer on the bottom of the well is analyzed and quantified (Fig. 1b, c). Due to the small size of the beads (typically less than 120 μm in diameter) a large number of technical repetitions is enabled and quantification with high statistical power is achieved (see Methods for details).

**Figure 1:**
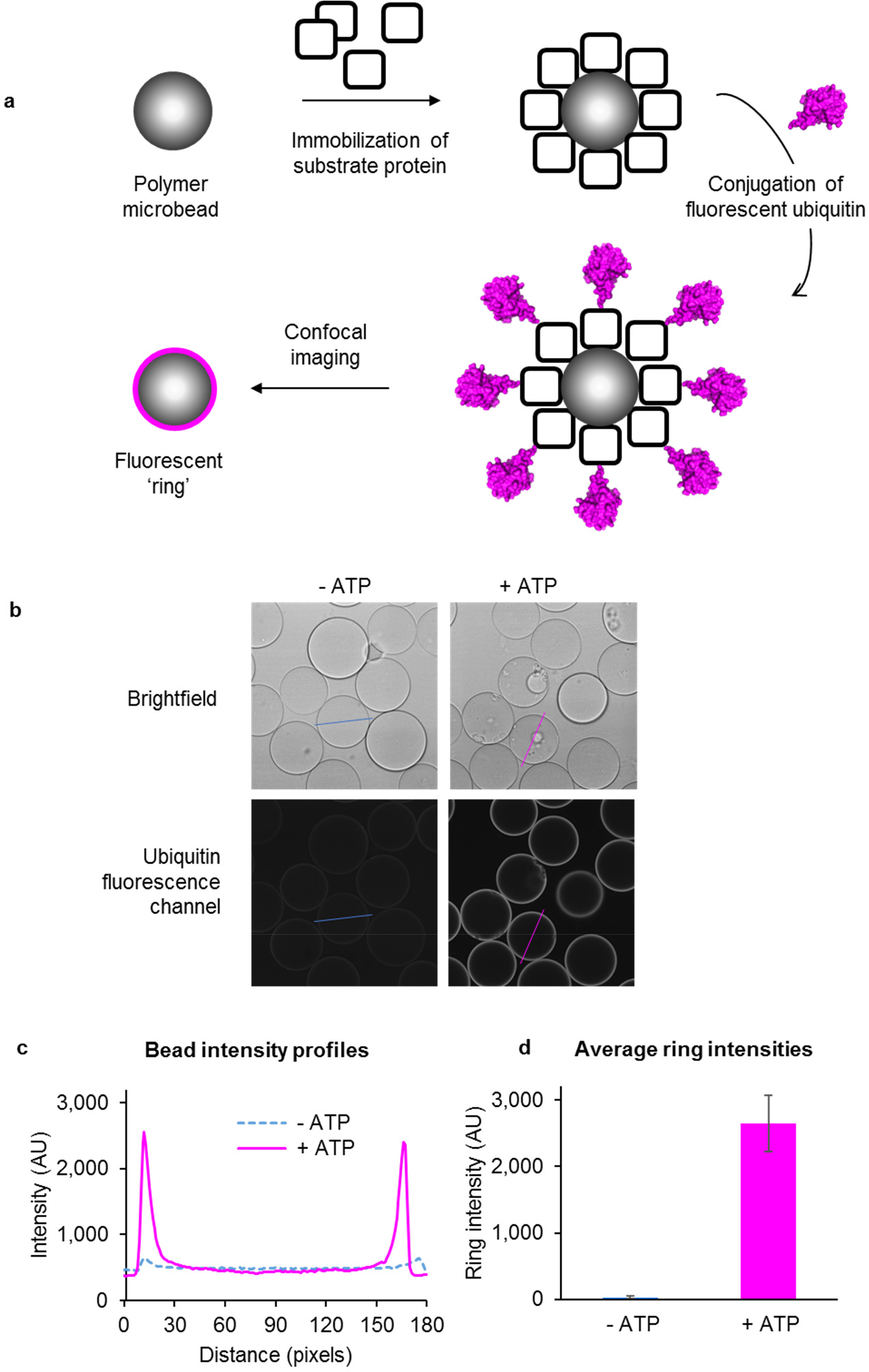
Detection of ubiquitination using Confocal Nanoscanning (CONA). (**a**) A substrate protein is immobilized on polymer microbeads and incubated with a suitable mix of conjugating enzymes and with a fluorescently labelled ubiquitin. Upon ubiquitin conjugation to the protein substrate, a fluorescent ‘ring’ will become detectable in the confocal imaging plane across the microbeads. (**b**) Images of beads in a 384-well plate were acquired using the confocal scanning microscope Opera™ (Perkin Elmer) in brightfield for bead detection (top images) and in the fluorescence emission channel of the dye on ubiquitin for detection of ubiquitin conjugates (bottom images). A reaction without ATP (-ATP) was used as control for possible non-enzymatic binding. Intensity profiles across beads (blue and magenta lines) were used for quantification of fluorescence emission intensity, which is proportional to the amount of conjugated ubiquitin at the concentrations used. (**c**) Bead cross-section intensity profiles from (**b**) were acquired using ImageJ software. (**d**) Intensity profiles of > 100 beads in a test well were analyzed as detailed in Methods to calculate average ring intensities in each well.

The on-bead ubiquitination reaction allows for detection of ubiquitin and ULM conjugates on a variety of enzymes and substrates (Fig. 2). The charging of ubiquitin to an E1 enzyme can be assayed by immobilizing the E1 on beads and incubating with ubiquitin in solution, together with Mg^2+^-ATP in reaction buffer. Alternatively, an E2 enzyme can be placed on bead and incubated with both the ubiquitin and an E1 enzyme in solution to detect thioester formation on the E2. Similarly, those E3 enzymes that form a direct thioester conjugate with ubiquitin, such as the HECT-type enzymes ^12^, can be tested by immobilizing the E3 and incubating with ubiquitin, E1 and an appropriate E2 in solution. Finally, the performance of an entire cascade can be quantified by immobilization of a substrate of interest on beads in the presence of ubiquitin and the appropriate E1, E2 and E3 enzymes in solution (Supporting Information Figure S. 2). Importantly, the flexibility, modularity and single-bead miniaturization of the UPS-CONA assay make it suitable for a plethora of various ubiquitin-related reactions of high complexity, which are difficult to investigate by existing methods. Using the same principle, the assay setup can also be applied to monitor the release of ubiquitin from an immobilized E1 into the E2 in solution, to track ubiquitination cascades in a step-by-step fashion (Fig. 2) or to simultaneously compare activities of enzymes from the same class.

**Figure 2:**
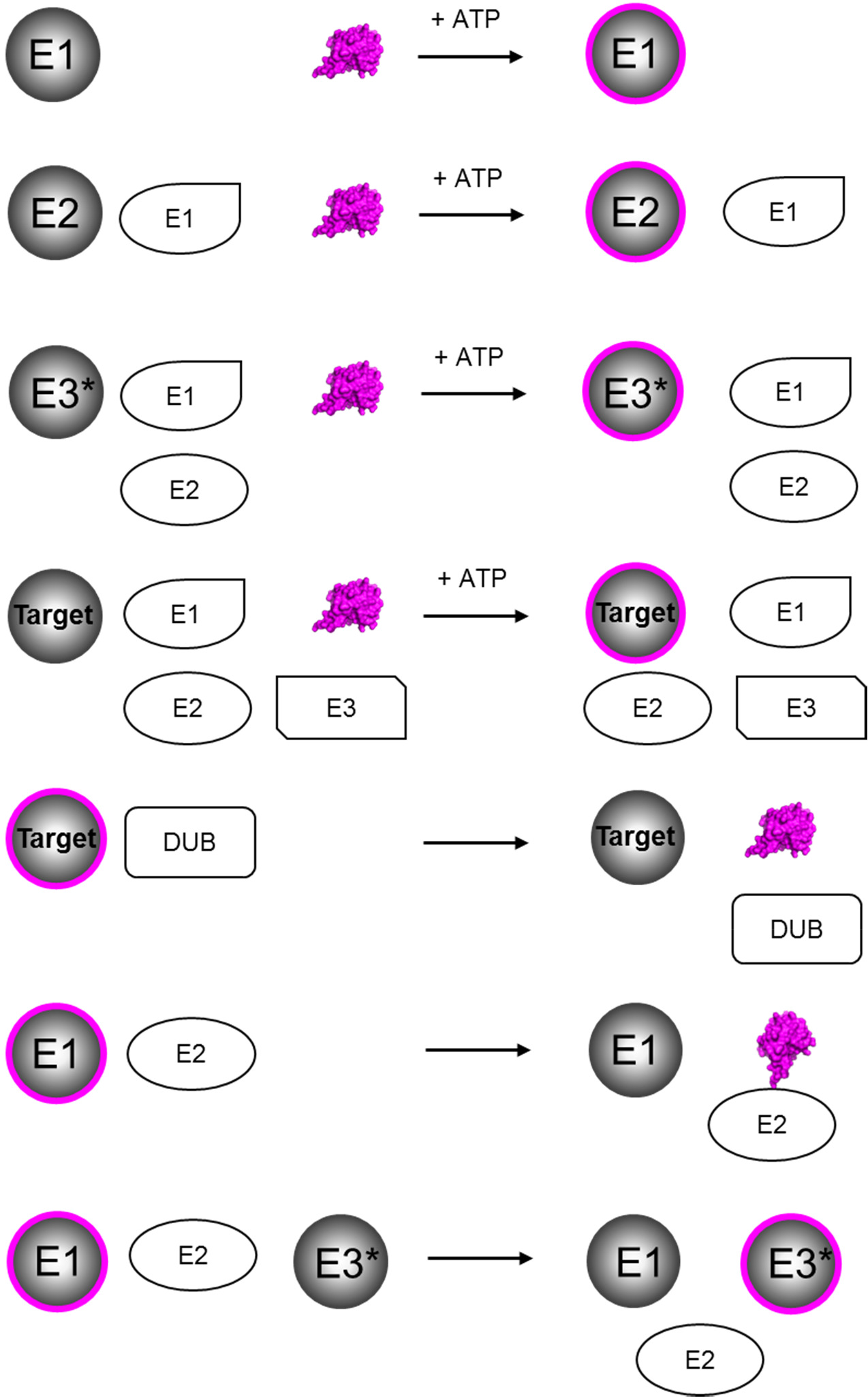
Applications of the UPS-CONA assay. UPS-CONA assay can be used to monitor various ubiquitin-related reactions. For example, ubiquitin conjugation to E1, E2 enzymes, HECT E3 ligase (E3*) and substrate proteins can be followed by immobilizing the proteins or enzymes of interest on polymer microbeads and incubating them with appropriate reaction components and with a fluorescently labelled ubiquitin. Increase in bead fluorescence will indicate a successful reaction. Ubiquitin release from an on-bead substrate and transfer to a downstream cascade component can also be detected as a decrease of bead fluorescence.

### Monitoring activities of E1, E2 and HECT E3 enzymes

To test the broad applicability of the UPS-CONA, we applied the method to different classes of ubiquitination enzymes, namely E1, E2 and HECT E3 enzymes. First, a His6-tagged version of the Ube1 ubiquitin-activating enzyme (E1) was immobilized on commercial Ni^2+^NTA agarose beads, previously sieved to limit their diameters to between 100 μm and 120 μm. The Ube1 beads were transferred to a 384-well plate to form a monolayer, and were incubated with Cy5-labelled ubiquitin (Cy5-Ub) in ubiquitination buffer as described in Methods. Blank beads were used as control for unspecific Cy5-Ub binding and reaction trials without ATP were used as a control for non-enzymatic Cy5-Ub binding to Ube1. Ubiquitin loading onto Ube1 was monitored by confocal detection of the Cy5 fluorescence emission on bead and quantified as detailed in the Methods section. The fluorescent emission intensity denoting ubiquitin conjugation increased rapidly, within minutes of incubation (Fig. 3a). Similar experiments were performed with an immobilized E2 (Ube2L3, Fig. 3b), in the presence of Cy5-Ub and an untagged Ube1 enzyme in solution. Finally, an immobilized version of the E6-AP ubiquitin ligase, which is a host cell factor for human papillomavirus E6-oncoprotein-induced degradation of p53 ^13^, was incubated with Cy5-Ub, untagged Ube1 and Ube2L3 in solution (E6AP, Fig. 3c). In all types of experiments, an increase of the Cy5 fluorescence intensity corresponding to ubiquitin thioester formation and/or conjugation was observed over time. A slight decrease of the signal observed after incubation for 90 minutes or longer was likely due to thioester hydrolysis. Addition of the reducing reagent dithiothreitol (DTT) at the end of each reaction revealed the proportion of conjugates attached through a thioester to the immobilized proteins to be: 60% on Ube1, 95% on Ube2L3 and 80% on E6AP (Fig, 3 a, b, c, respectively). The remaining signal on Ube1 is most likely to correspond to the ubiquitin adenylate bound to the adenylation site ^14^, while the remaining signal on E6AP likely corresponded to isopeptide bond-linked ubiquitin moieties due to auto-ubiquitination on E3 lysine residues ^15^. As expected, Ube2L3 was exclusively charged with thioester-bound ubiquitin under these conditions ^16^. We successfully tested several other ubiquitination and neddylation reactions using UPS-CONA, including ubiquitination of the tumor suppressor protein p53 (Supporting Information Table 1). Based on these results, the bead-based assay appears suitable for monitoring various reactions in the primary ubiquitination cascade, and in addition can be used to quantify the thioester-bound ubiquitin fraction relative to total bound ubiquitin.

**Figure 3:**
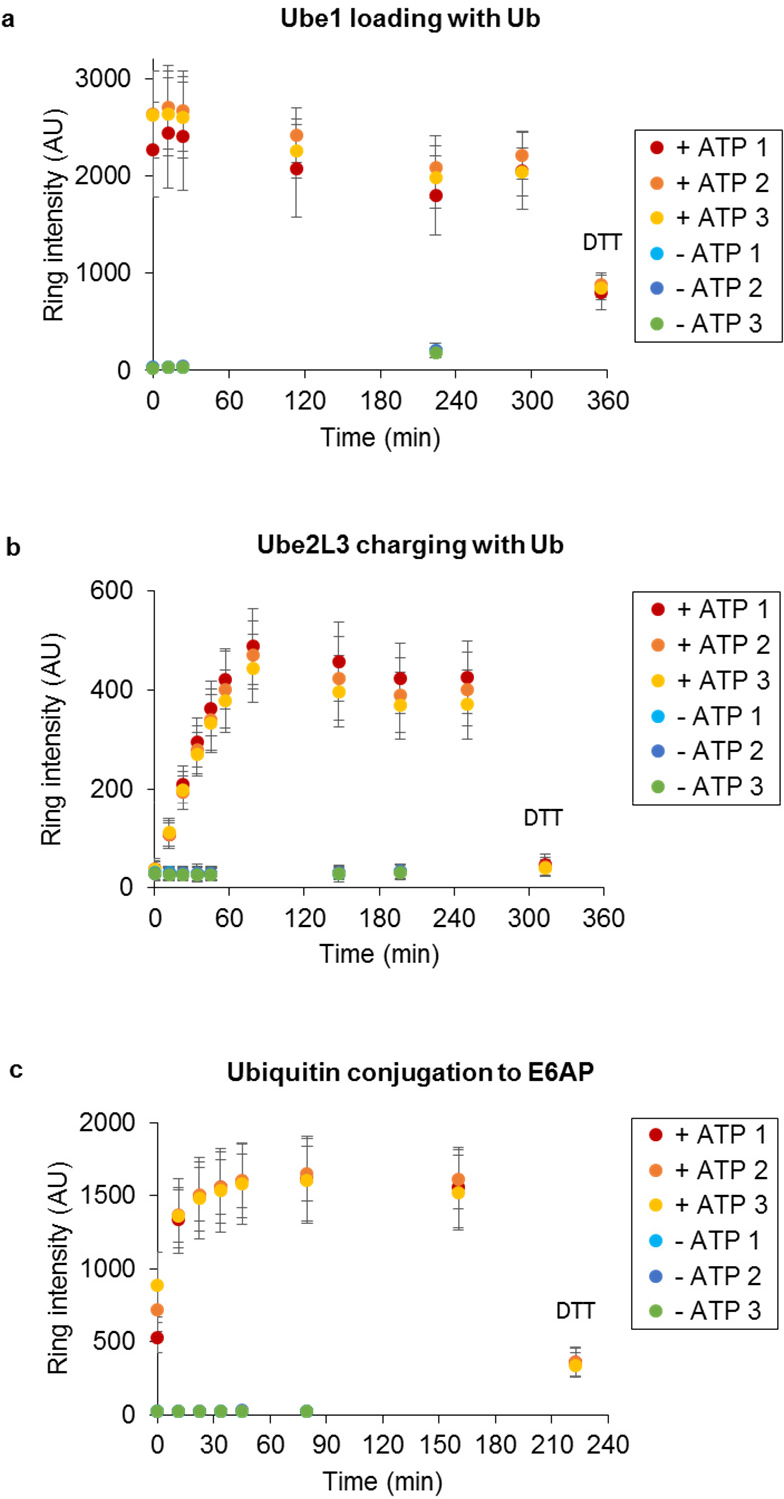
Ubiquitination activity of E1, E2 and HECT E3 enzymes detected over time using UPS-CONA. His6-tagged enzymes Ube1 (**a**), Ube2L3 (**b**) or E6AP (**c**) were immobilized on Ni^2+^NTA agarose microbeads, placed in a 384-well plate and incubated in the reaction buffer at 37°C in the presence of Cy5-labelled ubiquitin (**a**), Cy5-labelled ubiquitin and untagged Ube1 (**b**) or Cy5-labelled ubiquitin, untagged Ube1 and Ube2L3 (**c**). Reactions without ATP (-ATP) were used as controls for non-enzymatic binding of Cy5-Ub to the beads. Reactions were performed in triplicates. At the end of each reaction, DTT was added to remove thioester-bound Cy5-Ub from the on-bead enzymes to quantify non-thioester-bound fraction. Images were acquired on the confocal scanning microscope Opera™ (Perkin Elmer), in brightfield for detection of beads and in Cy5 fluorescence emission for detection of Cy5-Ub conjugates. Data were analyzed as described in the Methods. Average ring intensities in each well over time are represented, corresponding to Cy5-Ub conjugated to the on-bead enzyme.

### Multiplexing bead populations in a single well

Miniaturization of the assay, and subsequently a potential screening setup, to the single bead level, allows for modularity, multiplexing and flexibility in assay format. Each of the 200 - 400 beads placed into one well of a 384 well plate represents a separate technical repeat of the experiment, which thereby provides enhanced statistical power through multiple replicates per well.

To establish a system for investigation of multiple reactions, typical for enzymatic ubiquitination cascades, it was necessary to separately detect, within a well, bead populations to which different proteins were attached. The detection of mixed bead populations can be achieved through labelling of molecules involved in the reaction with spectrally distinct fluorophores and by then mixing these bead populations in a test well for detection with different fluorescence filters (Fig. 4a, b and Supporting Information Figure S 4). However, as this approach involves pre-labelling of proteins of interest with dyes, which could affect their activity, a method based on differentiating the bead populations by size was also established. Two bead populations were used: small beads of 40-70 μm in diameter and large beads of 100-120 μm in diameter (Fig. 4c). To confirm that the His6-tag-Ni^2+^NTA attachment was stable enough to prevent transfer between different bead populations in one well over time, small beads pre-incubated with His6-eGFP were mixed with blank large beads and vice versa in the same well (Fig. 4d). The small and large bead populations were separately detected without significant overlap by precisely setting the bead size parameters (Supporting Information Figure S 5). No protein exchange between the beads was observed under reaction conditions, likely due to local rebinding effects, which leads to strong compartmentalization of the His6-tagged proteins on the bead surface, in conjunction with non-saturating protein amounts (Supporting Information Figure S 1). These results demonstrated that each bead can be considered as a separate reaction unit, thereby allowing monitoring of more than one reaction in a single well.

**Figure 4:**
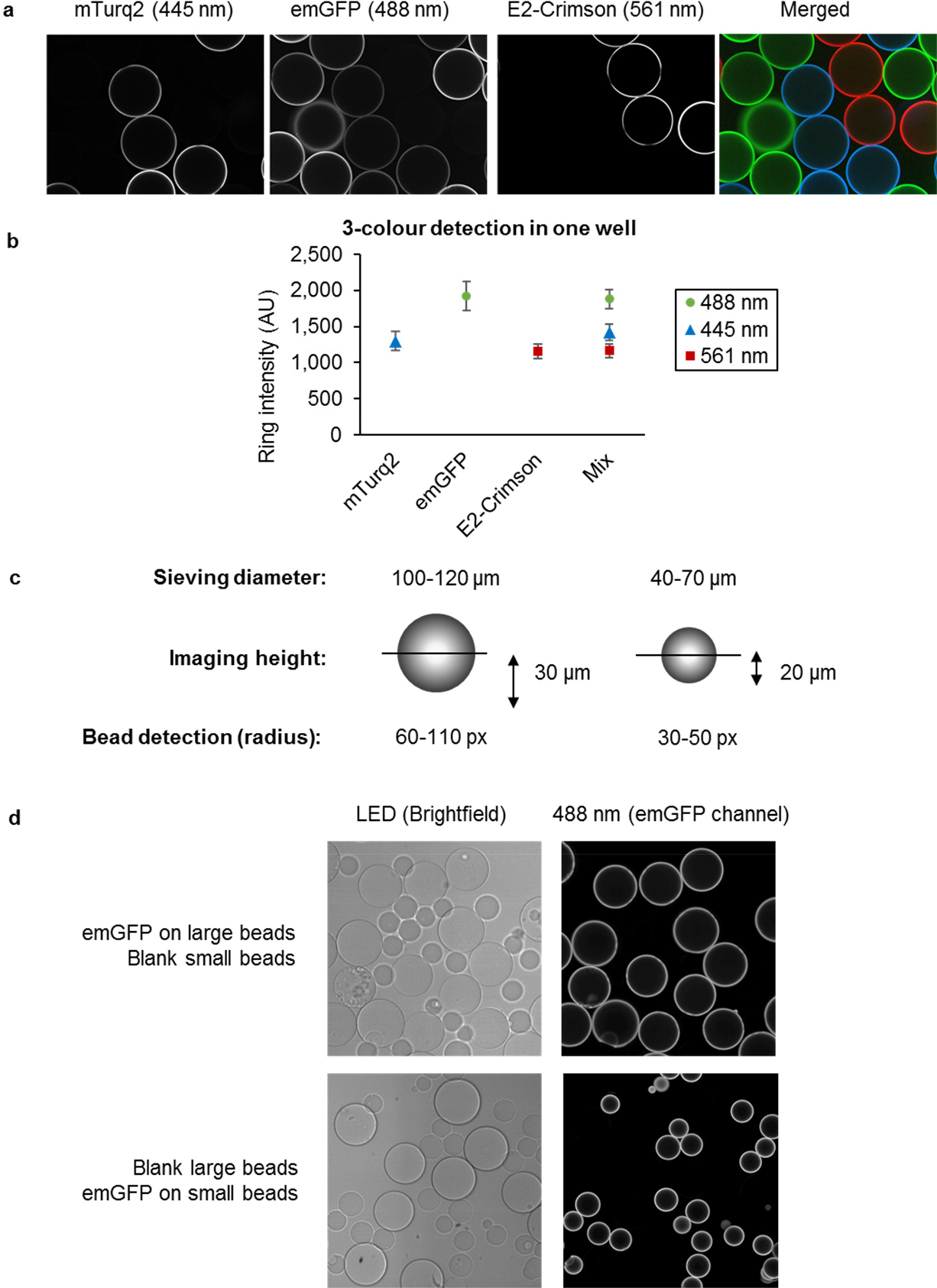
Detection of mixed bead populations by different color or by bead size.(a, b) Three bead-bound fluorescent proteins detected in a single well. (**a**) Three different His6-tagged fluorescent proteins (mTurquoise2 (mTurq2, blue), emGFP (green) and E2-Crimson (red)) were pre-immobilized on individual batches of Ni^2+^NTA agarose beads. The beads were then pooled together in one well of a 384-well plate. Images were acquired using the confocal scanning microscope Opera™ (Perkin Elmer) with 445 nm, 488 nm and 561 nm excitation wavelengths, and detected in three emission channels (475/34, 520/35, 660/150 and 690/70 nm, detailed in Supporting Information Figure S4). (**b**) Images of wells with single and mixed bead populations of mTurquoise2, emGFP and E2-Crimson beads were analyzed and average ring intensities were calculated for each channel. (**c**) **Differential detection of bead populations by size.** Large beads were sieved through pore size mesh of 100 and 120 μm, imaged on the Opera™ (Perkin Elmer) at a confocal plane height of 30 μm, and identified in the 60-110 pixel radius range. Small beads were sieved through pore size filters of 40 and 70 μm, imaged on the Opera at the confocal plane of 20 μm and detected in the 30-50 pixel radius range. (**d**) **Each bead is a separate reaction unit**. His6-tagged emGFP was immobilized on large or small beads and mixed in one well with blank small or blank large beads. Beads were detected as specified in (**c**) in brightfield and emGFP fluorescence emission channels. No emGFP was detected on the blank beads present in the same well, indicating the absence of protein transfer between the beads.

### Tracking a multi-step ubiquitination cascade

A challenge inherent to multi-enzyme reactions such as for the ubiquitination cascade, is the tracking of activities of various enzymes in a single experiment. To demonstrate the suitability of UPS-CONA for monitoring several serial steps in a reaction cascade, we chose to reconstitute the entire reaction for the E6AP, a clinically relevant E3 enzyme implicated in cervical cancer ^17^. A His6-tagged Ube1 enzyme was attached to small beads and a His6-tagged E6AP to large beads (Fig. 5a). The two bead populations were then mixed together in one well and incubated with the ubiquitination reaction solution, containing Cy5-Ub and ATP (see Methods). The small Ube1 beads became fluorescent (Fig. 5b) and the ring intensity reached a plateau within minutes, indicating a successful Cy5-Ub conjugation to Ube1. At this point, we added a non-tagged Ube2L3, the cognate E2 enzyme for E6AP. As the ubiquitin was transferred from the on-bead Ube1 to Ube2L3 in solution, the Cy5-Ub signal on Ube1-bound beads decreased over time (Fig. 5b, c). Simultaneously, the Cy5-Ub signal increased on the E6AP-bound large bead population, indicating that ubiquitin was conjugated from Ube2L3 in solution to the E6AP on beads (Fig. 5b, c). This result demonstrated the ability of the UPS-CONA assay to monitor multiple reactions in parallel in a single reaction well.

### Target deconvolution of ubiquitination inhibitors

Despite the therapeutic potential of the UPS in cancer and other disease areas, the activity and selectivity of many reported molecules that modulate the UPS are not fully characterized. In particular, assessment of the specificity of a potential inhibitor towards more than one reaction in a cascade requires development of multiple assays. We reasoned that an ideal assay system for addressing target specificity in the UPS should allow enzymatic reactions of the entire cascade to be monitored in parallel. Quantitative and time-resolved monitoring of ubiquitin transfer through the E1-E2-E3 cascade to the final substrate would provide mechanistic insight into the potential effects of an inhibitor at each step. This information would allow triage of hit compounds that lacked the necessary specificity for lead development. To investigate whether UPS-CONA can be used to assess selectivity of ubiquitination inhibitors, we used our multi-step ubiquitination assay to test a selection of known ubiquitination inhibitors: an E1 inhibitor PYR-41 ^18^, the E2 inhibitors BAY 11-7082 ^19^ and CC0651 ^20,21^, and the HECT E3 inhibitors clomipramine ^22^ and heclin ^23^.

**Figure 5:**
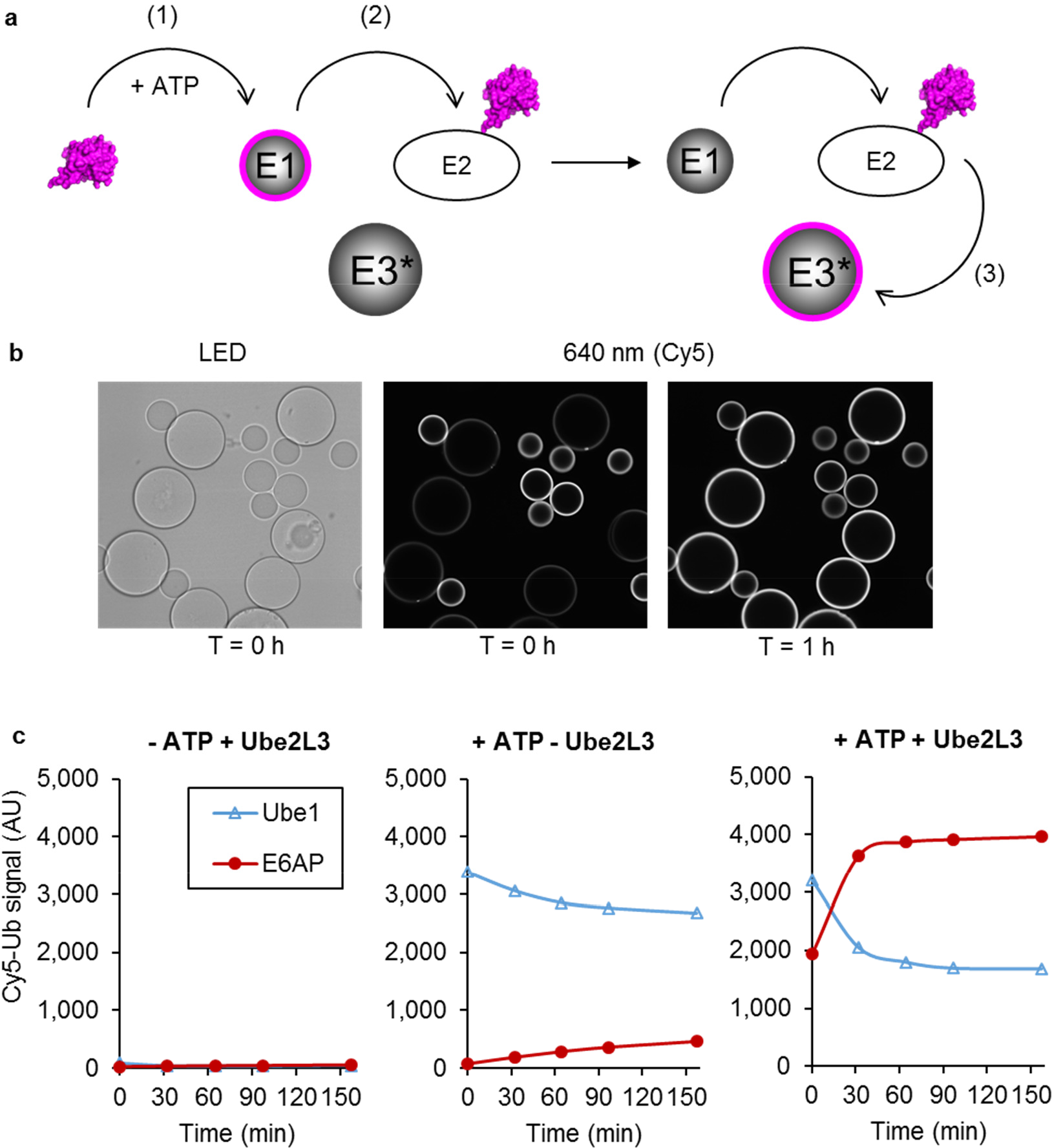
Real-time tracking of a ubiquitination cascade in a single well. (**a**) Schematic of the multi-step ubiquitination assay. Ube1 immobilized on small beads and E6AP immobilized on large beads were placed together in one well with a fluorescent Ub. In the presence of ATP, ubiquitin was loaded onto Ube1 (1), forming a fluorescent ring on small beads. Upon addition of Ube2L3 to the solution, ubiquitin was transferred to Ube2L3 and a decrease in ring intensity on small beads (Ube1) was observed (2). Simultaneously, as Ube2L3 further transfers the fluorescent Ub to E6AP, ring intensity on large beads increased (3). (**b**) The reaction was performed at 37°C and monitored by imaging on the confocal scanning microscope Opera™ (Perkin Elmer) over time. Images were acquired in the brightfield and Cy5 fluorescence emission detection channels. Shown are exemplary images at the start of experiment (T = 0 h) and after one hour of incubation (T = 1 h). Over time, the intensity of the Cy5-Ub conjugates decreased on Ube1 beads and increased on the E6AP beads. (**c**) Ring intensities from small (Ube1) and large (E6AP) beads were analyzed as described in Methods. Ubiquitin loading on Ube1 (blue line) and conjugation on E6AP (red line) which occurred simultaneously in each well are represented in charts, corresponding to wells as indicated: without ATP (− ATP + Ube2L3), without Ube2L3 (+ ATP − Ube2L3), with ATP and Ube2L3 (+ ATP + Ube2L3).

### BAY 11-7082 inhibits Ube1

The small molecule BAY 11-7082 was first identified as an inflammation inhibitor ^24^ and has been extensively used as such. More recently, Ube2L3 and other E2 enzymes and ubiquitination enzymes have been identified as direct targets of BAY 11-7082 ^19^. In our multi-activity UPS-CONA assay with Ube1, Ube2L3 and E6AP, we found that BAY 11-7082 decreased the rate of ubiquitin transfer for all individual reaction steps of the cascade (Fig. 6a). The combined, overall inhibitory effect of BAY 11-7082 on E6AP was stronger than that of the E1 inhibitor PYR-41 (Fig. 6a). To confirm the activity of BAY 11-7082 directly on Ube1, we tested the compound in an on-bead assay with Ube1 alone in the absence of E2 or E3 enzymes (Fig. 6b). The inhibitory effect of BAY 11-7082 on Ube1 was concentration-dependent, with an IC_50_ around 3 μM. To confirm that the observed inhibitory activity was not an artefact caused by effects on the fluorescent labels, we tested the compound in a standard SDS-PAGE ubiquitination assay. After optimizing the reaction conditions (see Supporting Methods), we determined that BAY 11-7082 inhibited Ube1 with an IC_50_ of ~4.6 μM in the gel-based assay (Fig. 6c). The magnitude of this effect was similar to the inhibition observed for other targets of BAY 11-7082 ^19^ and may in part explain the pan-inhibitory effect of BAY 11-7082 on E2 enzymes. In a similar experimental design, we observed that the E3 inhibitor heclin also affected ubiquitin loading onto Ube1 (Fig. 6b, d). The IC_50_ of 20 μM for this inhibitory effect of heclin was somewhat higher than the IC_50_ values of 6-7 μM reported for inhibition of other HECT E3s ^23^ but nonetheless suggested that E1 inhibition may also account for some activity of heclin. In contrast, we found that CC0651 and clomipramine had no detectable effect on Ube1 activity (Fig. 6b, d).

**Figure 6:**
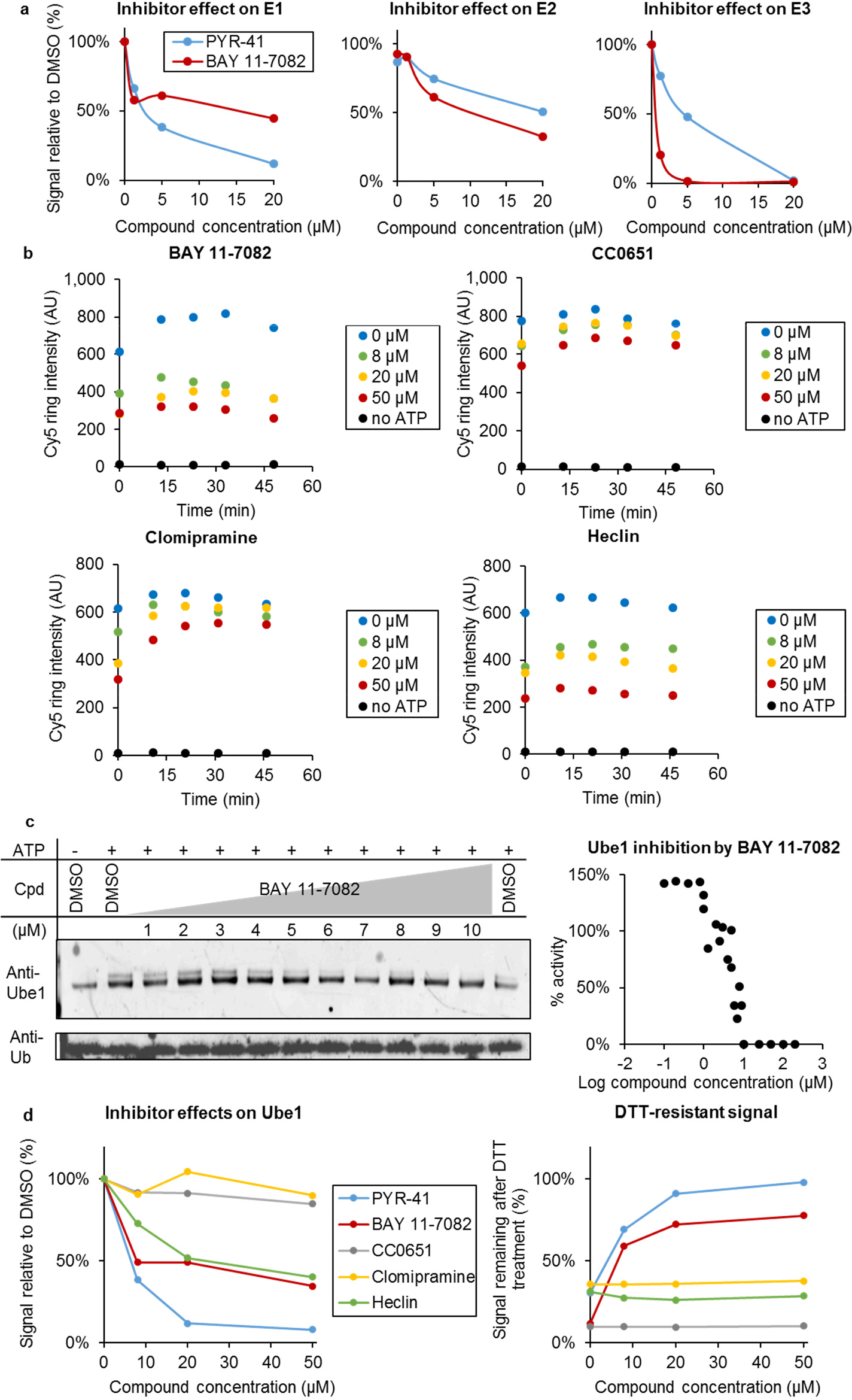
Characterization of ubiquitination inhibitors using UPS-CONA. BAY 11-7082 (pan-E2 inhibitor), CC0651 (Cdc34 inhibitor), clomipramine, heclin (HECT E3 inhibitors) and PYR-41 (Ube1 inhibitor) were tested for selectivity in the multi-step UPS-CONA assay. (**a)** Effects of BAY 11-7082 and PYR-41 on the Ube1-Ube2L3-E6AP cascade. Inhibition of Ube1 activity was measured as decrease in fluorescent ubiquitination signal on Ube1 prior to adding Ube2L3 to the reaction. Inhibition of Ube2L3 activity was measured indirectly as a decrease of Ub transfer from Ube1 to Ube2L3. Final effect on E6AP ubiquitination was measured as a decrease in end-point ubiquitin conjugation levels to E6AP. BAY 11-7082 inhibited the cascade already in the first step, namely Cy5-Ub conjugation to Ube1. (**b**) Detecting inhibitor effects on Ube1 activity using time-resolved UPS-CONA. Ube1 was immobilized on beads, preincubated with compounds at indicated concentrations for 15 minutes at 25°C and incubated with Cy5-Ub and ATP in ubiquitination buffer to measure inhibition effect. Images were acquired on the Opera™ instrument and analyzed to quantify ubiquitin loading on Ube1. BAY 11-7082 and heclin inhibited the reaction in a concentration-dependent manner. (**c**) Gel-based assay for BAY 11-7082 inhibition of Ube1. Ube1 was preincubated in the ubiquitination buffer with the test compound at indicated concentrations prior to adding WT-ubiquitin and ATP. After 5 minutes at 30°C, the reactions were stopped by adding SDS loading buffer. Samples were resolved by SDS-PAGE and revealed by Western blot with anti-ubiquitin and anti-Ube1 antibodies. Ube1 activity (Ube1~Ub thioester ratio to total Ube1) was quantified using densitometric analysis (ImageJ). The chart shows corresponding inhibition curve generated in four independent experiments. (**d**) Comparison of inhibition effects on Ube1 observed using UPS-CONA. On the left, dose-dependent inhibition of ubiquitin loading to immobilized Ube1 was observed for PYR-41, BAY 11-7082 and heclin. On the right, quantification of the non-thioester-bound ubiquitin fraction remaining after treatment with DTT revealed potentially different mechanisms of action of BAY 11-7082 and heclin.

### Mechanistic insights into enzyme inhibition

As the UPS-CONA readout combines quantitative and time-resolved detection of ubiquitination, we reasoned that it should be possible to resolve selected aspects of inhibition mechanisms. We therefore analyzed the dose-response effects of the aforementioned inhibitors on ubiquitin loading on Ube1 on beads, followed by reduction with DTT (Fig. 6d). In this format, the signal remaining after DTT treatment would correspond to ubiquitin that interacted with Ube1 through a thioester-independent mechanism. Analysis of the inhibition curves revealed that BAY 11-7082 inhibited up to ~50% of ubiquitin loading onto Ube1 (Fig. 6d, left), which suggested that this compound may act on only one ubiquitin binding site of Ube1. Since ubiquitin cannot occupy the second thioester site on Ube1 without binding first to the adenylation site, we inferred that the thioester site was likely affected. The DTT-resistant signal was dose-dependent, reaching ~80% at highest compound concentration used (Fig. 6d, right). This result indicated that the proportion of the thioester-bound ubiquitin was reduced by BAY 11-7082, consistent with an effect on the thioester site on Ube1. This mechanism implied a similar mode of action of BAY 11-7082 as described previously for E2 enzymes, by forming a covalent adduct with the reactive cysteine residues ^19^. In contrast, the effect of heclin on Ube1 was less pronounced, and similar to PYR-41 in the shape of the inhibition curve (Fig. 6d, left). However, the proportion of DTT-resistant signal remained unchanged upon increasing heclin concentrations (Fig. 6d, right), suggesting that heclin acts through a different mechanism than PYR-41 or BAY 11-7082 ^23^. As revealed by dose-response curves and DTT treatment, CC0651 and clomipramine did not affect Ube1 activity, which was confirmed in SDS-PAGE ubiquitination assays.

The multi-step monitoring capability of the UPS-CONA assay allowed for identification of a new inhibitory activity of BAY 11-7082 towards Ube1. This information is critical for the ubiquitin community, as this activity will need to be taken into account in assays using this compound so far known as a broad E2 enzyme, but not E1 enzyme inhibitor ^19^. To make decisions on further development of any hit compound from a binder screen or an enzymatic screen, the selectivity information is critically important. UPS-CONA resolves biochemical mode of action in one well of a 384-well plate with picomole amounts of reagents, and may be used for selectivity assays, either based on enzyme or ubiquitin mutations, or structure-activity relationships of hit series. Given the small amounts of enzymes with conventional tags needed for UPS-CONA, combined with the sensitivity and time resolved capabilities demonstrated in this work, this method should facilitate the study of reaction mechanisms, mode of action of inhibitors and high-throughput screening campaigns.

### Conclusion

We have developed and validated an on-bead confocal detection method called UPS-CONA which is suitable for studying individual enzymatic activities, including various E1, E2 and HECT-type E3 enzymes. We also show that the various ubiquitin-E1-E2-E3 reaction intermediates of a ubiquitination cascade can be followed in a single reaction vessel using an exemplary enzymatic cascade from Ube1 through Ube2L3 to E6AP in a time-resolved manner. A dynamic, confocal detection of fluorescent ubiquitin conjugates ensures a timely, sensitive readout, and a straightforward protocol can be easily automatized for large-scale experiments. Furthermore, our integrated assay is suitable for studying effects of small molecule inhibitors upstream or downstream of the presumptive target against which an inhibitor was originally identified.

UPS-CONA may be parallelized further using different bead sizes to allow simultaneous monitoring of ubiquitination of different substrates in the same reaction or adapted to monitor deubiquitinating enzyme and ubiquitin binding domain activities. Ubiquitin-like modifiers, for example SUMO or NEDD8, may also be monitored in similar assay formats or used for bead-based deconvolution of multiple modification types on a single substrate. Given the robustness of the on-bead fluorescent ring readout to interfering fluorescence from the reactions solutions ^7,8^, a lysate-based assay may also be used to study reactions under more physiological conditions, without the need for prior purification of reaction components. The suitability of off-the-shelf confocal imaging instrumentation and commercially available reagents should enable UPS-CONA technology to become a broadly applicable method. UPS-CONA should therefore be of value for exploring the interactions and activities within combinatorially complex UPS cascades, for identification and characterization of new small molecule probes, and for facilitating UPS drug discovery.

## METHODS

### Bead preparation

Nickel nitrilotriacetic acid (Ni^2+^NTA) agarose micro-beads were purchased from Qiagen (cat. n° 30250). Before use, beads were filtered through various mesh sizes (40, 70, 100 and 120
μm) using cell strainers (Corning cat. 352340, 352350 and 352360) or nylon woven net filters (Millipore cat. n° NY2H04700) to obtain beads in specific size ranges. Beads were then washed thoroughly with binding buffer (0.3 M NaCl, 20 mM HEPES pH 7.5, 0.01% Triton X-100) and brought back to 50% slurry. 1 μL of 100-120 μm filtered beads per well of a 384-well plate were used to form a monolayer with around 50% coverage of the well bottom surface. The volumes and amounts were scaled up according to the required number of wells.

### Immobilizing protein on beads

The estimated bead loading capacity as provided by the manufacturer is around 50 mg per 1 mL of the 50 % slurry solution for a protein of around 30 kDa. 1 μL of beads would have a protein loading amount equal to 50 μg of protein. Generally, we used a maximum of 0.5 μM in 20 μL of on-bead protein per well corresponding to 300 ng, which is over 160-fold below saturating amounts. The optimal amount of protein on bead for effective signal detection in our system was determined by using His6-emGFP. Below an amount of 20 pmoles/well we detected a linear relationship between the amount of on-bead emGFP and ring intensity (Supporting Information Figure S1). We also verified that the His6-tagged protein was stably attached to the beads for several hours which is probably due to fast on and off rates corresponding to μM dissociation constants given by the tagging system (Supporting Information Figure S3a). The beads were incubated with the desired quantity of His6-tagged protein (typically 2.5 pmoles per well, or 125 nM) in ice-cold binding buffer on a shaker at 1000 rpm for at least 30 min at 8°C. After incubation, the beads were extensively washed with binding buffer and the volume of bead solution was adjusted to 10 μL per well.

### On-bead ubiquitination reaction

Beads with attached proteins were distributed in 10 μL volumes into microplate wells (black, flat glass bottom 384-well plate, MMI PS384B-G175) using a wide bore pipette tip (Rainin RC-250W). Ubiquitination mixes were prepared to obtain final concentrations in 20 μL as follows: 500 nM Cy5-labelled ubiquitin (Cy5-Ub), 50 nM Ube1, 1 μM Ube2L3, 5 mM ATP where appropriate, in 50 mM Tris pH 7.5, 5 mM MgCl_2_. Ubiquitin-activating reactions with on-bead Ube1 were performed with Cy5-Ub and ATP. Ubiquitin-charging reactions with on-bead Ube2L3 were performed with Cy5-Ub, Ube1 and ATP. Ubiquitin-charging/conjugation to on-bead E6AP were performed with Cy5-Ub, Ube1, Ube2L3 and ATP. The plate was incubated at 37°C and the reactions were monitored over time. For determination of proportion of thioester-bound ubiquitin, 50 mM DTT was added at the end of the reaction and imaged again after 10 min incubation.

### Bead imaging

Images were acquired on the Opera™ High Content Screening System (Perkin Elmer) at 30.0 μm above well bottom, at 20 × magnification with air lenses (20X Air LUCPLFLN, NA=0.45). Brightfield and two fluorescent channels for detection of emGFP and Cy5 were used in the following settings: excitation wavelength 640 nm (Cy5), 488 nm (emGFP); emission filters: 690/70 nm (Cy5), 520/35 nm (emGFP). Typically, 77 images from the centre of the well were taken to ensure near-full well coverage and to visualize >100 beads per well, and a well sublayout with 20% image field overlap was applied to allow consequent image stitching. Image data were analyzed using ImageJ and a custom MatLab script.

### Data analysis

Full well images are stitched together from multiple individual microscope images using the Grid/Collection Stitching Plugin for ImageJ ^25^. The stitched images from each channel, either brightfield or fluorescent, are analyzed for each well in a well plate. The detection in the brightfield channel allows detection of all the beads in a given well, including those with only weak or no fluorescence signal. An edge detection algorithm and Gaussian filtering is applied to the image, followed by a circular Hough transfer to identify circles in the image within a required diameter range (typically between 100 and 120 μm) representing the beads. A canonical list of bead positions and radii in each well is compiled by correlating the beads detected from all the imaging channels. Beads which are incomplete, overlapping or coincide with the image edges are removed from the list. A pre-determined number of profiles (typically 10) is taken across each bead. The peaks representing the ‘ring’ and the mean of the 20-80th percentiles of the profile intensity between the peaks representing the background bead intensity are identified for each profile. A profile ‘intensity’ is computed by subtracting the background from the peak intensity. The mean of these background-corrected ring intensity measurements for each bead is computed, giving the bead intensity. If more than one fluorophore is used in the experiment, the ratio of fluorescence emission intensities of two fluorophores detected on bead is used for an enhanced quantification as it allows for correction against a reference fluorophore, such as a second label on the assayed protein immobilized on beads. Ratiometric analysis is performed by calculating the ratio of the bead intensity of the same bead in multiple fluorescent channels. The mean bead ring intensity of each well in each channel is calculated as the average bead intensity of all the beads in the well. For each well, the mean bead ring intensity and standard deviation are calculated. The optimal amount of protein on bead for effective signal detection was determined by using a His6-tagged Emerald Green Fluorescent Protein (emGFP) standard. A linear correlation between the amount of on-bead emGFP and ring intensity held for values equal to and below 10 pmoles/well (Supporting Information Figure S1). It was also verified that His6-tagged proteins were stably attached to the beads for several hours under typical experimental conditions (Supporting Information Figure S3).

## Acknowledgements

This work was supported by the Wellcome Trust PhD Programme grant n° 089397/Z/09/Z. M.A. also acknowledges financial support from the Scottish Universities Life Sciences Alliance ((SULSA-http://www.sulsa.ac.uk), the Medical Research Council ((MRC-www.mrc.ac.uk, J54359) Strategic Grant, and from the European Community’s 7th Framework Program (FP7/2007–2013) under grant agreement no 278568 ‘PRIMES’. F.S. and M.T. acknowledge support from the Canadian Institutes for Health Research (MOP-126129). The funders had no role in study design, data collection and analysis, decision to publish, or preparation of the manuscript.

## Supporting Information

**Supporting Methods and Supporting Information Figures S1-S6**

## Author Contributions

J.K. co-invented and developed the assay, designed, performed and analyzed the experiments and wrote the manuscript; N.P. contributed to confocal microscopy experiments, produced labelled ubiquitin and participated in data analysis; D.E. created the software for data analysis and supported analysis of the results; S.M. cloned the fluorescent proteins into expression vectors; I.P. helped with data analysis; M.T. co-invented the assay; D.C. and F.S. provided plasmids and protein reagents and supported with ubiquitin purification and labelling; S.S. contributed to data analysis and to figure generation, M.A. developed the concept of expanding on-bead screening to enzymatic reactions exemplified in this work with UPS-CONA, co-invented the assay, and wrote the manuscript.

## Competing Financial Interests

The University Court of the University of Edinburgh filed an international patent application n° PCT/GB2015/052565 which covers parts of the method.

## References

1. Petroski, M. D. The ubiquitin system, disease, and drug discovery. BMC Biochem. 9 Suppl 1, S7 (2008).

2. Huang, X. & Dixit, V. M. Drugging the undruggables: exploring the ubiquitin system for drug development. Cell Res. 26, 484–498 (2016).

3. Goldenberg, S. J., Marblestone, J. G., Mattern, M. R. & Nicholson, B. Strategies for the identification of ubiquitin ligase inhibitors. Biochem. Soc. Trans.38, 132–6 (2010).

4. Berkers, C. R. & Ovaa, H. Drug discovery and assay development in the ubiquitin-proteasome system. Biochem. Soc. Trans. 38, 14–20 (2010).

5. Eldridge, A. G. & O’Brien, T. Therapeutic strategies within the ubiquitin proteasome system. Cell Death Differ. 17, 4–13 (2010).

6. Hintersteiner, M. et al. Confocal nanoscanning, bead picking (CONA): PickoScreen microscopes for automated and quantitative screening of one-bead one-compound libraries. J. Comb. Chem. 11, 886–94 (2009).

7. Hintersteiner, M. et al. Single bead labeling method for combining confocal fluorescence on-bead screening and solution validation of tagged one-bead one-compound libraries. Chem. Biol. 16, 724–35 (2009).

8. Hintersteiner, M., Buehler, C. & Auer, M. On-bead screens sample narrower affinity ranges of protein-ligand interactions compared to equivalent solution assays. Chemphyschem13, 3472–80 (2012).

9. Meisner, N.-C. et al. Terminal adenosyl transferase activity of posttranscriptional regulator HuR revealed by confocal on-bead screening. J. Mol. Biol. 386, 435–50 (2009).

10. Hintersteiner, M. et al. Identification of a small molecule inhibitor of importin β mediated nuclear import by confocal on-bead screening of tagged one-bead one-compound libraries. ACS Chem. Biol. 5, 967–79 (2010).

11. Hintersteiner, M. et al. Identification and X-ray co-crystal structure of a small-molecule activator of LFA-1-ICAM-1 binding. Angew. Chem. Int. Ed. Engl. 53, 4322–6 (2014).

12. Huibregtse, J. M., Scheffner, M., Beaudenon, S. & Howley, P. M. A family of proteins structurally and functionally related to the E6-AP ubiquitin-protein ligase. Proc. Natl. Acad. Sci. U. S. A. 92, 2563–7 (1995).

13. Scheffner, M., Huibregtse, J. M., Vierstra, R. D. & Howley, P. M. The HPV-16 E6 and E6-AP complex functions as a ubiquitin-protein ligase in the ubiquitination of p53. Cell75, 495–505 (1993).

14. Haas, A. L., Warms, J. V, Hershko, A. & Rose, I. A. Ubiquitin-activating enzyme. Mechanism and role in protein-ubiquitin conjugation. J. Biol. Chem. 257, 2543–8 (1982).

15. Nuber, U., Schwarz, S. E. & Scheffner, M. The ubiquitin-protein ligase E6-associated protein (E6-AP) serves as its own substrate. Eur. J. Biochem. 254, 643–9 (1998).

16. Nuber, U., Schwarz, S., Kaiser, P., Schneider, R. & Scheffner, M. Cloning of human ubiquitin-conjugating enzymes UbcH6 and UbcH7 (E2-F1) and characterization of their interaction with E6-AP and RSP5. J. Biol. Chem. 271, 2795–800 (1996).

17. Beaudenon, S. & Huibregtse, J. M. HPV E6, E6AP and cervical cancer. BMC Biochem. 9 Suppl 1, S4 (2008).

18. Yang, Y. et al. Inhibitors of ubiquitin-activating enzyme (E1), a new class of potential cancer therapeutics. Cancer Res. 67, 9472–81 (2007).

19. Strickson, S. et al. The anti-inflammatory drug BAY 11-7082 suppresses the MyD88-dependent signalling network by targeting the ubiquitin system. Biochem. J. 451, 427–37 (2013).

20. Ceccarelli, D. F. et al. An allosteric inhibitor of the human Cdc34 ubiquitin-conjugating enzyme. Cell145, 1075–87 (2011).

21. Huang, H. et al. E2 enzyme inhibition by stabilization of a low-affinity interface with ubiquitins. Nat. Chem. Biol. 10, 156–63 (2014).

22. Rossi, M. et al. High throughput screening for inhibitors of the HECT ubiquitin E3 ligase ITCH identifies antidepressant drugs as regulators of autophagy. Cell Death Dis. 5, e1203 (2014).

23. Mund, T., Lewis, M. J., Maslen, S. & Pelham, H. R. Peptide and small molecule inhibitors of HECT-type ubiquitin ligases. Proc. Natl. Acad. Sci. U. S. A. 111, 16736–41 (2014).

24. Pierce, J. W. et al. Novel Inhibitors of Cytokine-induced I B Phosphorylation and Endothelial Cell Adhesion Molecule Expression Show Anti-inflammatory Effects in Vivo. J. Biol. Chem. 272, 21096–21103 (1997).

25. Preibisch, S., Saalfeld, S. & Tomancak, P. Globally optimal stitching of tiled 3D microscopic image acquisitions. Bioinformatics25, 1463–5 (2009).

